# Genomic diversity analysis of SARS-CoV-2 genomes in Rwanda

**DOI:** 10.1101/2020.12.14.422793

**Authors:** Lambert Nzungize, Pacifique Ndishimye, Fathiah Zakham

## Abstract

COVID-19 (Coronavirus disease 2019) is an emerging pneumonia-like respiratory disease of humans and is recently spreading across the globe.

**Objective:** To analyze the genome sequence of SARS-CoV-2 (severe acute respiratory syndrome coronavirus-2) isolated from Rwanda with other viral strains from African countries.

**Methods:** We downloaded 75 genomes sequences of clinical SARS-CoV-2 from the GISAID (global initiative on sharing all influenza data) database and we comprehensively analyzed these SARS-CoV-2 genomes sequences alongside with Wuhan SARS-CoV-2 sequences as the reference strains.

**Results:** We analyzed 75 genomes sequences of SARS-CoV-2 isolated in different African countries including 10 samples of SARS-CoV-2 isolated in Rwanda between July and August 2020. The phylogenetic analysis of the genome sequence of SARS-CoV-2 revealed a strong identity with reference strains between 90-95%. We identified a missense mutation in four proteins including orf1ab polyprotein, NSP2, 2’-O-ribose methyltransferase and orf1a polyprotein. The most common changes in the base are C > T. We also found that all clinically SARS-CoV-2 isolated from Rwanda had genomes belonging to clade G and lineage B.1.

**Conclusions:** Tracking the genetic evolution of SARS-CoV-2 over time is important to understand viral evolution pathogenesis. These findings may help to implement public health measures in curbing COVID-19 in Rwanda.

## Introduction

COVID-19 disease caused by SARS-CoV-2 virus (Severe Acute Respiratory Syndrome Coronavirus-2) a divergent RNA (ribonucleic acid) virus for the present devasting the world, with over 60 million cases and more than 1.4 million deaths until November 25, 2020 (1). The SARS-CoV-2 genome replication and how COVID-19 disease spread rapidly in the early phase across the globe contributed to the genetic diversity inside the genome sequence of SARS-CoV-2 including clades (2, 3). Scientists isolated the virus and named SARS-CoV-2, they found a high nucleotide sequence homology with Betacorona viruses isolated from bats such as bat-SL-CoVZXC21 and bat-SL-CoVZXC21 (88%) and Malayan pangolins Pangolin-CoV (85.5% to 92.4%)(4). Interestingly, SARS-CoV-2 share similarity with other emerging human coronaviruses: SARS-CoV (79.5%) and the Middle East Respiratory Syndrome Coronavirus, MERS (50%) (5, 6).

The family of coronaviridae under the betacoronavirus genus as an RNA virus causes various diseases in birds and mammals, including SARS-CoV-2. It contains an RNA polymerase with proofreading activity(7, 8). The SARS-CoV-2 genome contains ∼29800 bp (9), therefore S-protein (spike protein) covers the surface of SARS-CoV-2, and consists of 1273 amino acids with a protein size between 180-200 kDa. The S-protein is composed of two subunits (S1 and S2) which play a key role in the cell membrane fusion process and receptor recognition, hence Spike protein becomes a target for vaccine development(10) (11, 12). The analysis of the SARS-CoV-2 genome sequence showed that Spike protein and RNA polymerase endure constant mutations according to geographic location (Asia, Europe and America). The SARS-CoV-2 genome analysis and generation of data have been a key component of discovering SARS-CoV-2 pathogenicity (13-15). Due to the frequent travels link between Rwanda to other geographic regions like China, United Arab Emirates, Europe, America and other African countries. It enhanced the chance to spot the first COVID-19 case on March 14, 2020; in Rwanda and the initial period of total lockdown started seven days later on March 21, 2020 (16). We hypothesized that the viral evolution of the SARS-CoV-2 is associated with the mutations within the SARS-CoV-2 genome. We aimed to track evolution and mutation within nucleotides sequence and proteins of SARS-CoV-2 genome by phylogenetic analysis to understand the pathogenesis and viral genome of SARS-CoV-2 in Rwanda.

## Methods

In total, we downloaded 75 genomes sequences of SARS-CoV-2 from the global initiative on sharing all influenza data (GISAID) (https://www.gisaid.org/) database, such as 10 genomes isolated in Rwanda, 3 genomes isolated in Code d’Ivoire, 2 genomes isolated in Zimbabwe, 1 genome isolated in Zambia, 4 genomes isolated in Uganda, 1 genome isolated in Tunisia, 3 genomes isolated in South Africa, 3 genomes isolated in Sierra Leone, 5 genomes isolated in Senegal, 3 genomes isolated in Reunion, 3 genomes isolated in Congo, 3 genomes isolated in Mozambique, 3 genomes isolated in Morocco, 2 genomes isolated in Madagascar, 3 genomes isolated in Kenya, 2 genomes isolated in Gambia, 2 genomes isolated in Gabon, 3 genomes isolated in Equatorial Guinea, 3 genomes isolated in Egypt, 3 genomes isolated in Democratic Republic of Congo, 3 genomes isolated in Algeria, 1 genome isolated in Cameroon, 2 genomes isolated in Botswana, 3 genomes isolated in Benin. These genomes sequences were analyzed with 4 Wuhan genomes isolated in China using the MEGA software version 10.0.4 with default parameters (17). All targeted genomes of SARS-CoV-2 samples were aligned to the reference strains using the Maximum Parsimony method by the Subtree-Pruning-Regrafting algorithm (18). The nucleotide variants within the SARS-CoV-2 genomes in the coding regions were performed by Genome Detective Coronavirus Typing Tool(19).

## Results

### Phylogenetic analysis

We compared the SARS-CoV-2 query sequences isolated in Rwanda from July to August 2020 to the other SARS-CoV-2 genomes isolated in Africa with Wuhan strains. The analysis of phylogenomic indicated that SARS-CoV-2 isolated in Rwanda belongs to lineage B.1. We found a significant sequence similarity of 90-95% between the WH-Human1_China_2019-Dec genome sequence and the SARS-CoV-2 present in Rwanda. Our data add to important information about the evolution of SARS-CoV-2 in Rwanda (Supplementary figure S1).

### Mutational analysis

The lineage assessment among the different SARS-CoV-2 lineages (A, B and B.1) indicated that lineage B.1 dominant in the SARS-CoV-2 genome isolated from Rwanda. All SARS-CoV-2 genomes sequence from Rwanda presents a universal type of mutations.

## DISCUSSION

The government of Rwanda implemented public health measures to reduce the spread of COVID-19 like national lockdown, social distance between people, wearing face masks in public. Therefore, the tracing showed that COVID-19 disease was maintained within the cities of Rwanda and no further spread of SARS-Cov-2 in the entire population, which resulted in a lower number of positive COVID-19 cases (16) (Fig. 1). The availability of the genome sequence of SARS-CoV-2 supports valuable resources to understand the viral evolution of SARS-CoV-2 (20).

**Figure 1.**
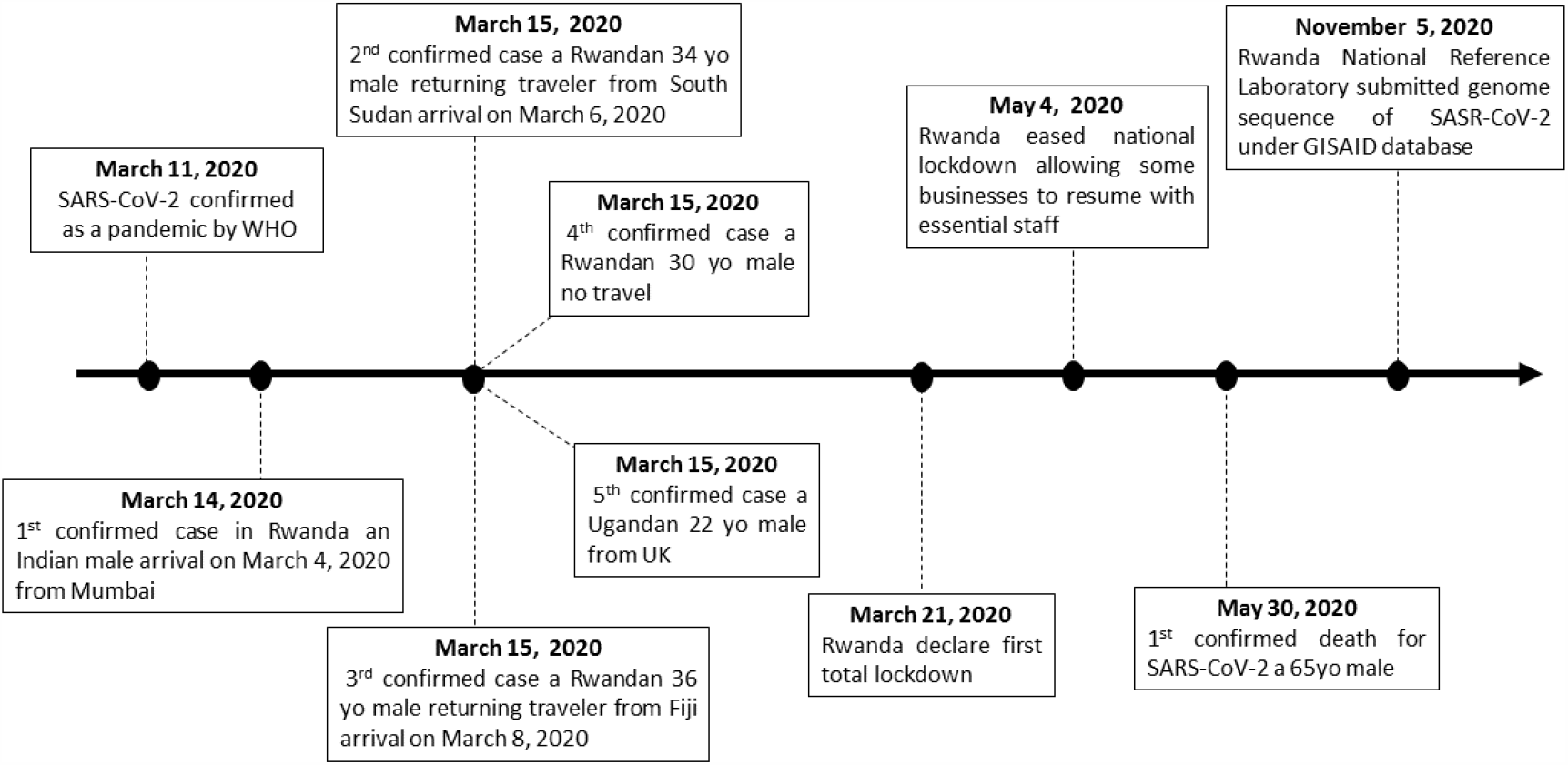
Timeline of COVID-19 and key events following the first confirmed case in Rwanda. (yo: years old).

**Figure 2.**
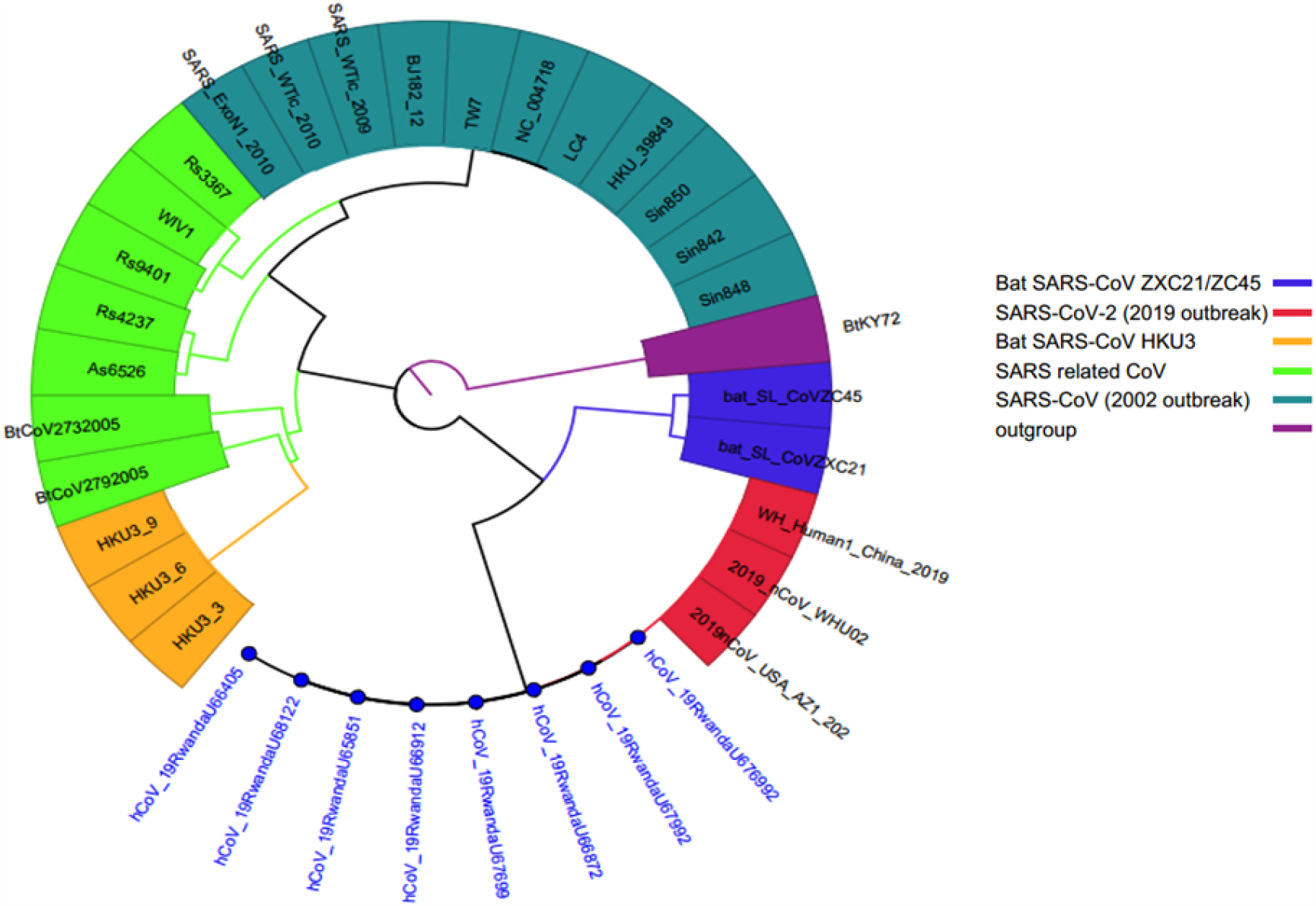
SARS-CoV-2 genomes sequences analysis from Rwandan patients. Bat SARS-CoV ZXC21/ZC45 (Accession Number: MG772934, MG772933.1) belong to SARSr-CoV Cluster isolated from China under the Bat animal. Bat SARS-CoV HKU3 (Accession Number: DQ084200, GQ153541, GQ153544) isolated in China under Bat animal (Table 2). The color corresponding to each sample within the same clade.

The genetic variation of the virus is classified into three clades (S, V, and G) (21); hence we note that clade G appears to be predominant in ten clinical sequences of SARS-CoV-2 isolated from Rwanda. The clade G was mostly spread in North America and Europe (22, 23). The phylogenetic analysis of the SARS-CoV-2 genome from Rwanda showed to be belonging to phylogenetic lineage B.1, which is sharing a common ancestor with the SARS-CoV-2 strain isolated in Europe, America, Asia and Australia (24). Further, the evolutional history was inferred using the Maximum Parsimony method with 0.223 of consistency index (CI), 0.636 of retention index (RI) and 0.142 of the composite index, therefore if CI is less than 0.5 designated that much homoplasies has occurred which is similar to our findings (25). The percentage of replicated tree taxa clustered indicated 90-95% (Supplementary figure S1). We performed a mutational analysis by observing changes in bases within SARS-CoV-2 genome (Table 3) and we found that all genomes sequence of SARS-CoV-2 isolated from Rwanda has a common type of mutations (Table 4) such as non-coding variant at position 241 (C to T) which affect 5’-UTR genome segment, 1746 (T to C), synonymous at position 3037 (C to T) which affect ORF1ab/NSP3 protein, 7420 (C to T), 12439 (C to T), missense at position 14408 (C to T), 22524 (A to T), missense at position 23403 (A to G) which affect Spike protein, 25186 (G to T), 26017 (T to G). The open reading frames (ORF10, ORF6, PRF3a, ORF 7a, ORF7b and PRF1ab) are predicted to encode for S-protein (S), NSP (non-structural proteins), (N) nucleocapsid proteins, envelope (E) and membrane (M) (26, 27). Although the SARS-CoV-2 genomes sequences used in this study indicated ten nucleotides difference compared to the reference strains that belong to G clade. Similar results of the type of mutations in SARS-CoV-2 were previously reported within the G clade (28). The genetic variations among the genome sequences of SARS-CoV-2 from Africa could be linked with the different geographical locations of each country.

**Table 1.** The sequence identity of beta coronavirus isolated in Rwanda used in this study.

| **GISAID**<br>**accession ID** | **Virus name** | **Location** | **Collection date** | **Age, Sex** |
| --- | --- | --- | --- | --- |
| EPI_ISL_615075 | Betacoronavirus | Rwanda | July, 2020 | 23, Male |
| EPI_ISL_615071 |  | Rwanda | July, 2020 | 23, Male |
| EPI_ISL_615069 |  | Rwanda | July, 2020 | 40, Male |
| EPI_ISL_615067 |  | Rwanda | July, 2020 | 58, Male |
| EPI_ISL_615066 |  | Rwanda | July, 2020 | 52, Male |
| EPI_ISL_615064 |  | Rwanda | July, 2020 | 30, Male |
| EPI_ISL_615063 |  | Rwanda | July, 2020 | 33, Male |
| EPI_ISL_614892 |  | Rwanda | July, 2020 | 40, Male |
| EPI_ISL_614891 |  | Rwanda | July, 2020 | 58, Male |
| EPI_ISL_614763 |  | Rwanda | August, 2020 | 52, Male |

**Table 2.**
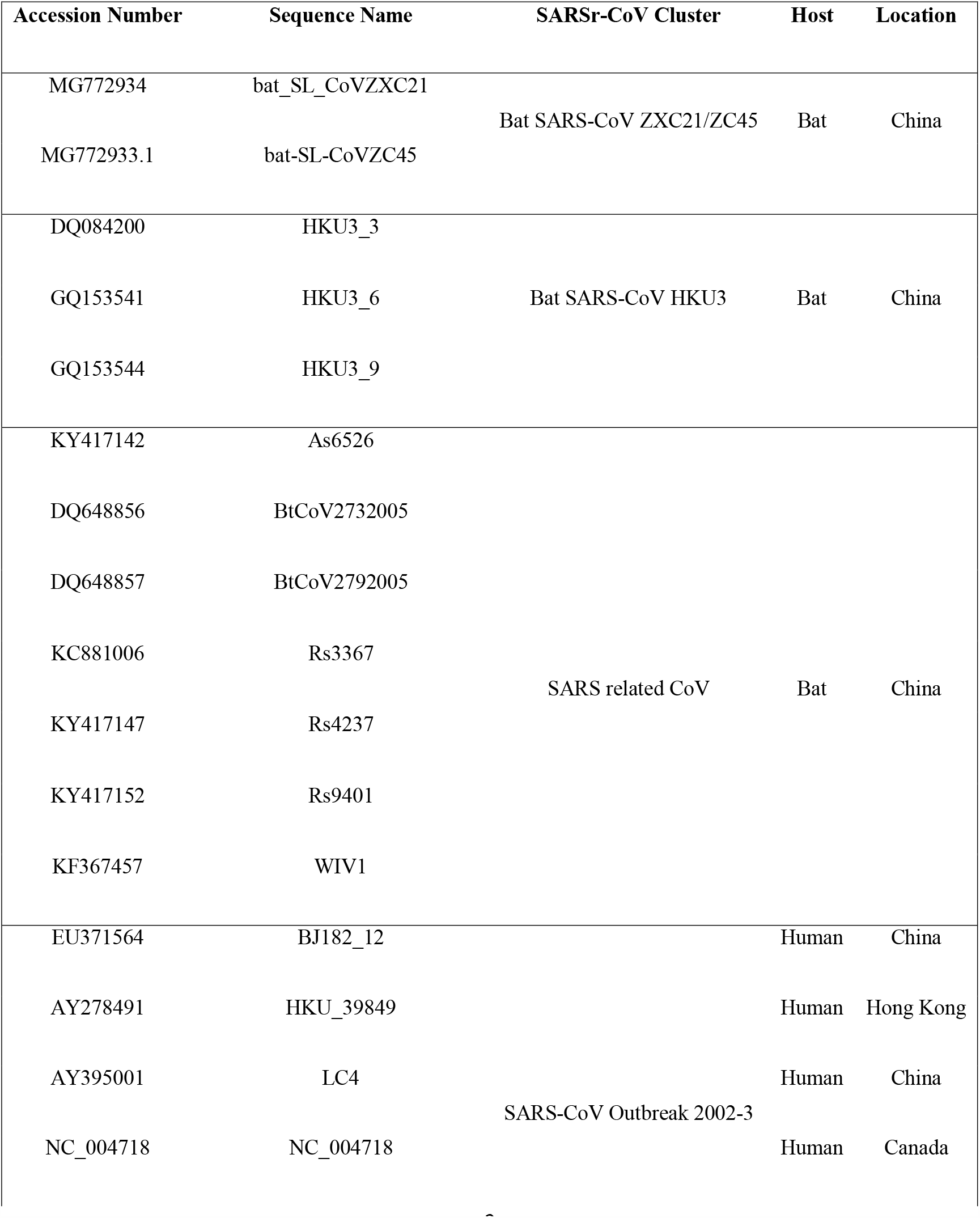
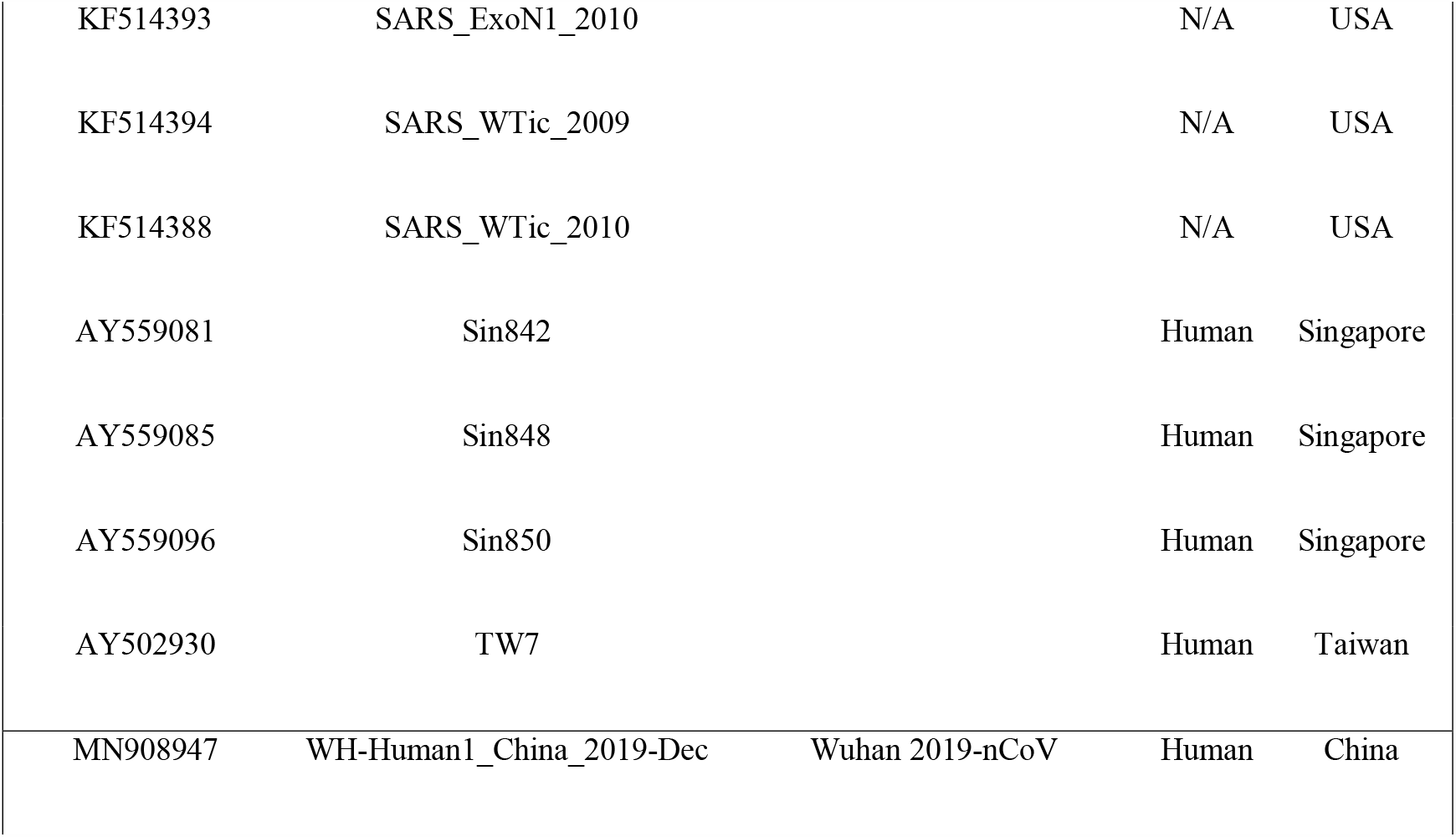
Reference cladogram dataset used to analyze genomes sequence of SARS-CoV-2 isolated in Rwanda.

| Accession Number | Sequence Name | SARSr-CoV Cluster | Host | Location |
| --- | --- | --- | --- | --- |
| MG772934 | bat_SL_CoVZXC21 | Bat SARS-CoV ZXC21/ZC45 | Bat | China |
| MG772933.1 | bat-SL-CoVZC45 |  |  |  |
| DQ084200 | HKU3_3 | Bat SARS-CoV HKU3 | Bat | China |
| GQ153541 | HKU3_6 |  |  |  |
| GQ153544 | HKU3_9 |  |  |  |
| KY417142 | As6526 | SARS related CoV | Bat | China |
| DQ648856 | BtCoV2732005 |  |  |  |
| DQ648857 | BtCoV2792005 |  |  |  |
| KC881006 | Rs3367 |  |  |  |
| KY417147 | Rs4237 |  |  |  |
| KY417152 | Rs9401 |  |  |  |
| KF367457 | WIV1 |  |  |  |
| EU371564 | BJ182_12 | SARS-CoV Outbreak 2002-3 | Human | China |
| AY278491 | HKU_39849 |  | Human | Hong Kong |
| AY395001 | LC4 |  | Human | China |
| NC_004718 | NC_004718 |  | Human | Canada |
| KF514393 | SARS_ExoN1_2010 |  | N/A | USA |
| KF514394 | SARS_WTic_2009 |  | N/A | USA |
| KF514388 | SARS_WTic_2010 |  | N/A | USA |
| AY559081 | Sin842 |  | Human | Singapore |
| AY559085 | Sin848 |  | Human | Singapore |
| AY559096 | Sin850 |  | Human | Singapore |
| AY502930 | TW7 |  | Human | Taiwan |
| MN908947 | WH-Human1_China_2019-Dec | Wuhan 2019-nCoV | Human | China |

**Table 3.** Nucleotide sequence and mutational analysis of the SARS-CoV-2 genome from Rwanda with the reference strain.

| Reference strain | Begin | End | Coverage | Score | Concordance | Matches | Identities | I/D/M/F | Stop<br>Codons |
| --- | --- | --- | --- | --- | --- | --- | --- | --- | --- |
| WH-Human1_China_2019-Dec | 55 | 29836 | 95.1% | 56842 | 99.9% | 28443 (100%) | 28431 (99.9%) | 0/0 |  |
| **CDS** |  |  |  |  |  |  |  |  |  |
| 1_orf1ab | 1 | 7097 | 95.1% | 47014 | 99.8% | 6749 (100%) | 6742 (99.9%) | 0/0/2/0 | 1 |
| 2_orf1a | 1 | 4406 | 96.2% | 29092 | 99.8% | 4237 (100%) | 4233 (99.9%) | 0/0/1/0 | 1 |
| 3_S | 1 | 1274 | 100% | 8961 | 99.7% | 1274 (100%) | 1271 (99.8%) | 0/0/0/0 | 1 |
| 4_ORF3a | 1 | 276 | 100% | 1952 | 99.8% | 276 (100%) | 275 (99.6%) | 0/0/0/0 | 1 |
| 5_E | 1 | 76 | 100% | 474 | 100% | 76 (100%) | 76 (100%) | 0/0/0/0 | 1 |
| 6_M | 1 | 223 | 100% | 1538 | 100% | 223 (100%) | 223 (100%) | 0/0/0/0 | 1 |
| 7_ORF6 | 1 | 62 | 100% | 400 | 100% | 62 (100%) | 62 (100%) | 0/0/0/0 | 1 |
| 8_ORF7a | 1 | 40 | 32.8% | 264 | 96.4% | 40 (100%) | 39 (97.5%) | 0/0/0/0 | 0 |
| 9_ORF7b | 19 | 44 | 59.1% | 195 | 98.0% | 26 (100%) | 26 (100%) | 0/0/0/0 | 1 |
| 10_ORF8 | 1 | 122 | 100% | 918 | 100% | 122 (100%) | 122 (100%) | 0/0/0/0 | 1 |
| 11_N | 1 | 420 | 100% | 2868 | 100% | 420 (100%) | 420 (100%) | 0/0/0/0 | 1 |
| 12_ORF9b | 1 | 98 | 100% | 618 | 100% | 98 (100%) | 98 (100%) | 0/0/0/0 | 1 |
| 13_ORF10 | 1 | 39 | 100% | 269 | 100% | 39 (100%) | 39 (100%) | 0/0/0/0 | 1 |
| 14_ORF14 | 1 | 74 | 100% | 502 | 100% | 74 (100%) | 74 (100%) | 0/0/0/0 | 1 |

**CDS:** coding sequencing; **Begin:** position of first nucleotide based on reference strain; **End:** position of last nucleotide that matches with reference strain; **Coverage:** coverage of reference strain genome; **Score:** nucleotide and amino acid score of AGA; **Matches:** number of matches; **Identities:** number of identical nucleotides; **I:** insertions; **D:** deletions; **M:** misaligned; **F:** Frameshifts; **Stop codons:** number of stop codons.

**Table 4.**
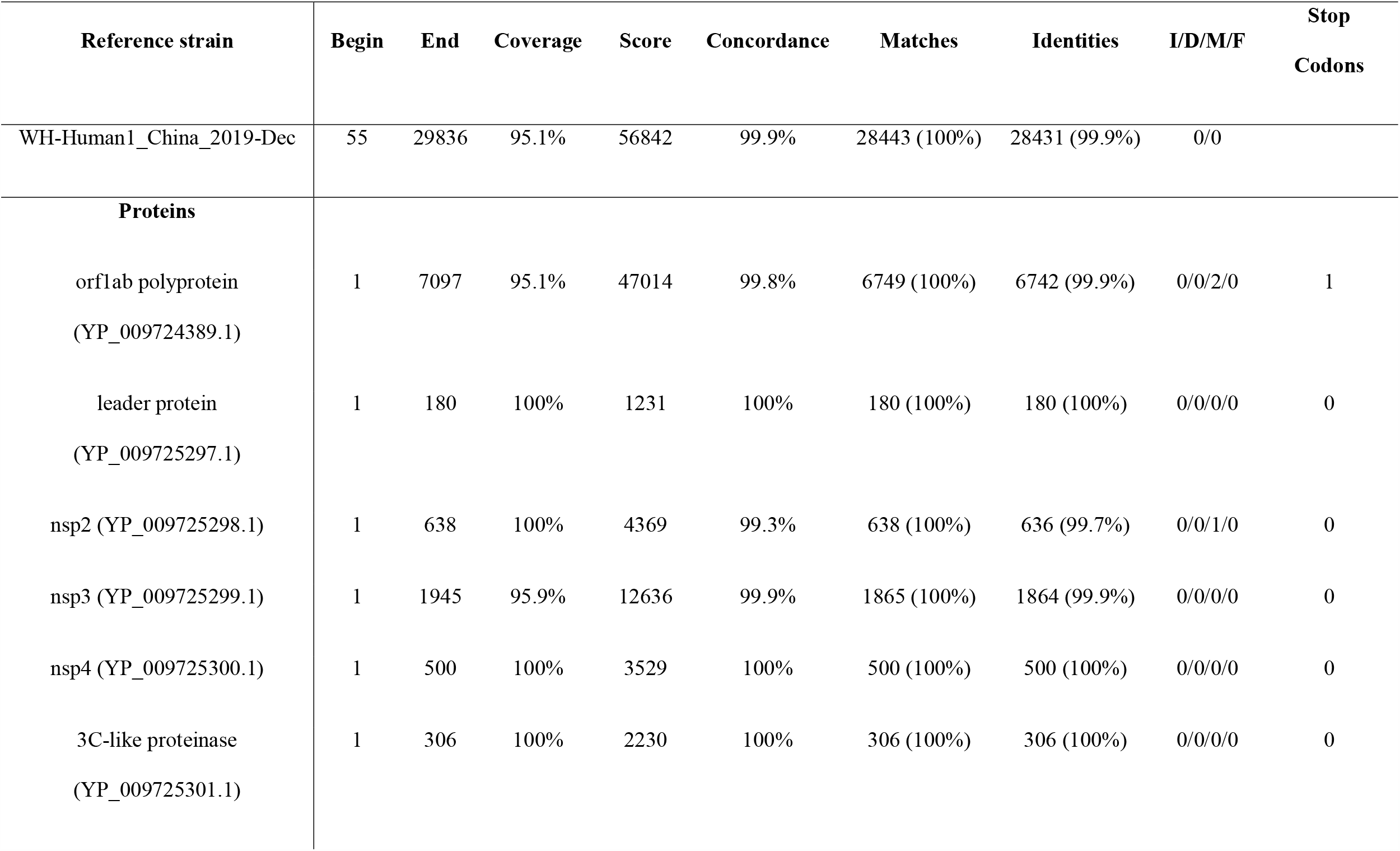
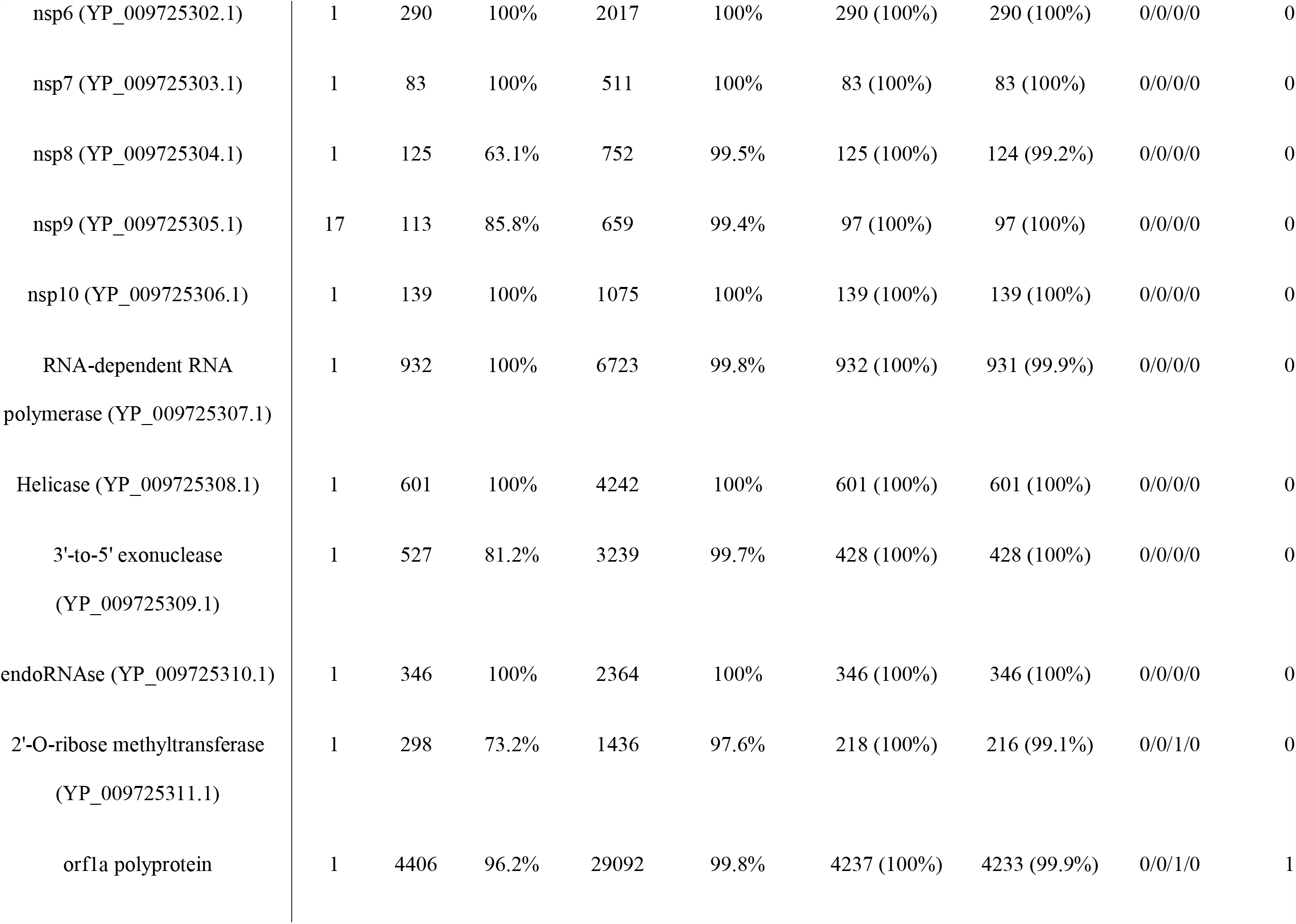
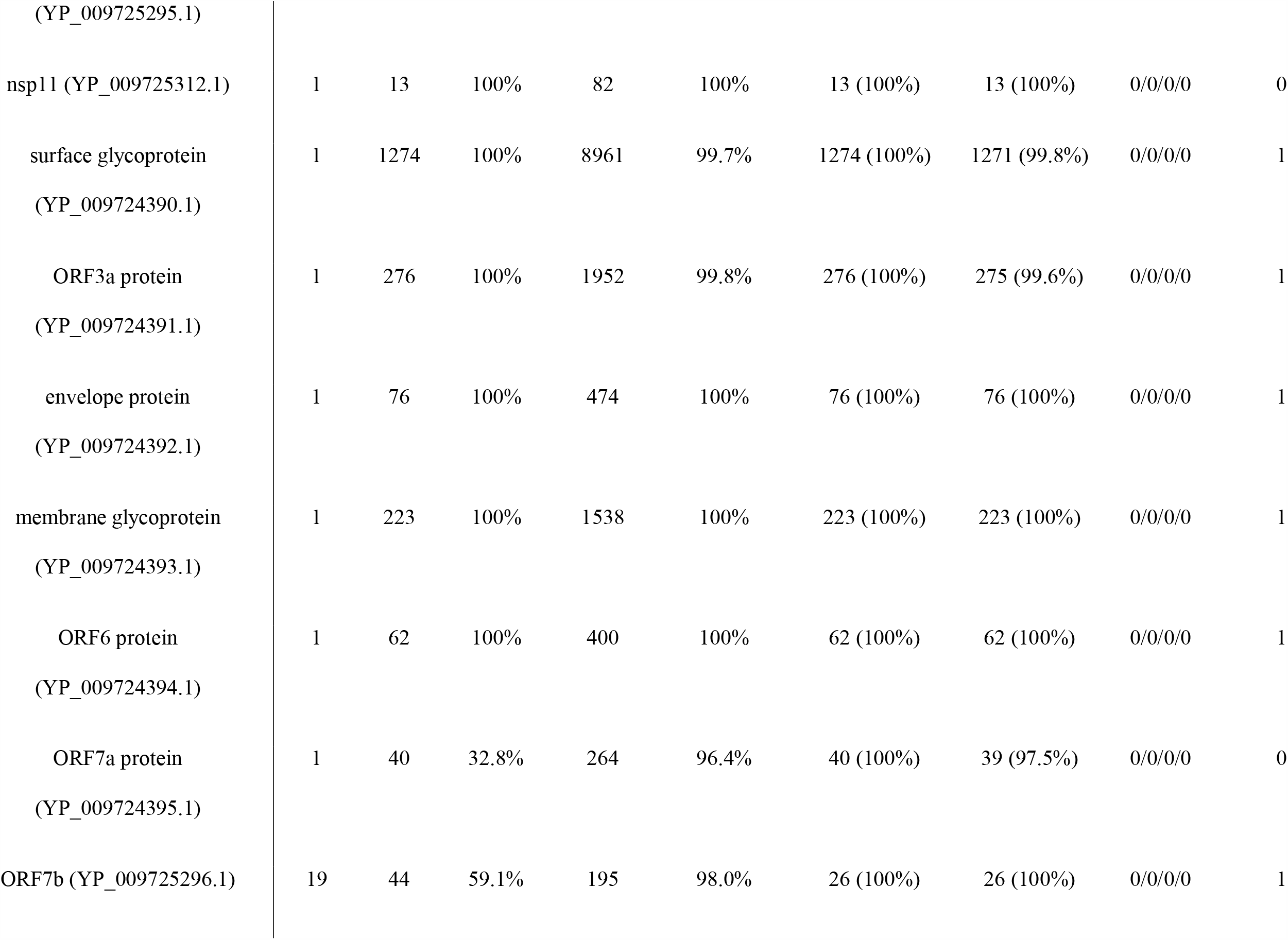
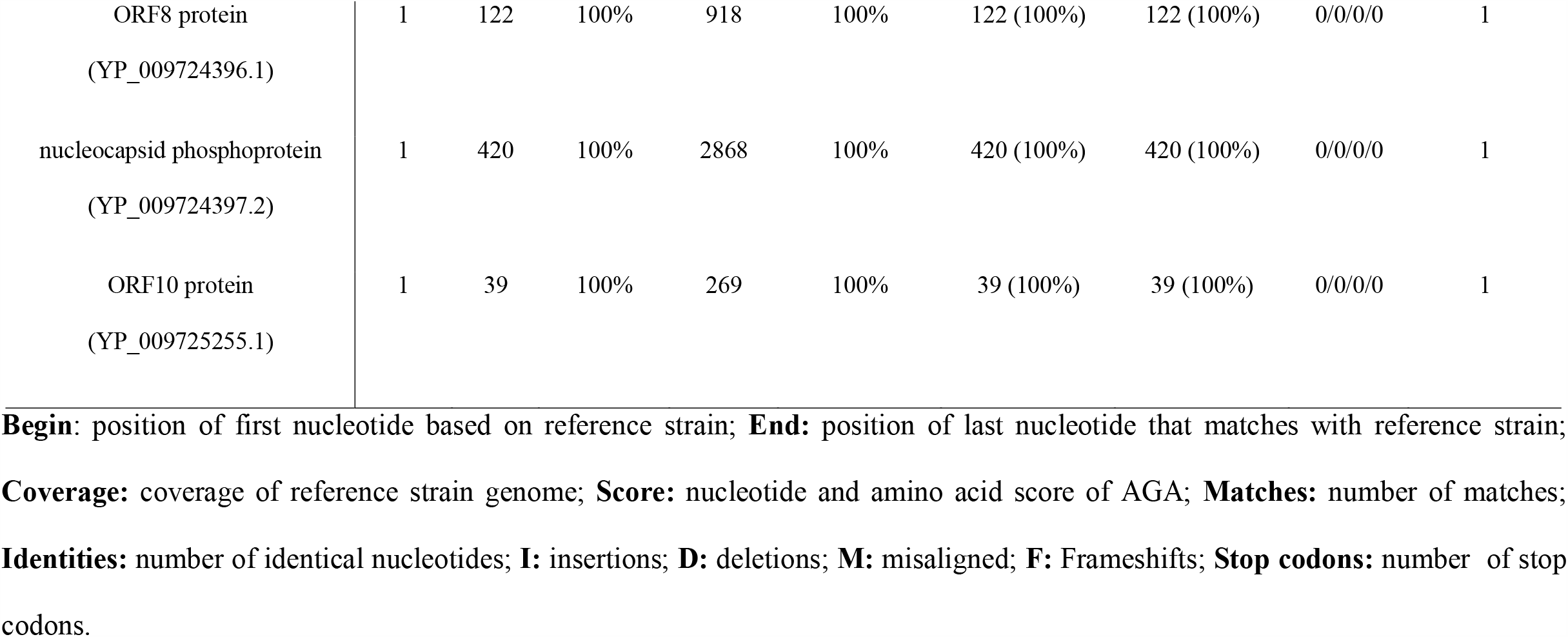
Proteins mutation of the SARS-CoV-2 from Rwanda compares to the reference strain

## Conclusion

The phylogenetic analysis shows genetic variation and the common genes or proteins but doesn’t rule out the probability that the importation of the SARS-CoV-2 virus originates from Europe, Asia or America. We recommend that the utility of epidemic surveillance based on genome sequence analysis as a key factor in monitoring the evolution and origin of SARS-CoV-2 in Rwanda.

## Authors contribution

All the author read and approved the final manuscript.

## Figure Legends

**Supplementary figure S1**. Phylogenic tree analysis of the SARS-CoV-2 genomes from Rwanda with other viral strains from African countries alongside with Wuhan strains. The tree was constructed by the Maximum Parsimony method with MEGA software (ver 10.0.4). The branch number indicates the percentage of bootstrap values and the scale represents branch length with reference strains.

## References

1. Worldometers. COVID-19 coronavirus pandemic 2020 [Available from: https://www.worldometers.info/coronavirus/?

2. V’Kovski P, Kratzel A, Steiner S, Stalder H, Thiel V. Coronavirus biology and replication: implications for SARS-CoV-2. Nature reviews Microbiology. 2020:1–16.

3. Lambert N, Umuhoza D, Dai Y, Nzaou SAE, Asaad M, Abokadoum M. Estimation of fatality rate in Africa through the behavior of COVID-19 in Italy relevance to age profiles. medRxiv. 2020.

4. Lam TT, Jia N, Zhang YW, Shum MH, Jiang JF, Zhu HC. Identifying SARS-CoV-2-related coronaviruses in Malayan pangolins. Nature. 2020;583(7815):282–5.

5. Lu R, Zhao X, Li J, Niu P, Yang B, Wu H. Genomic characterisation and epidemiology of 2019 novel coronavirus: implications for virus origins and receptor binding. Lancet (London, England). 2020;395(10224):565–74.

6. Wu F, Zhao S, Yu B, Chen YM, Wang W, Song ZG. A new coronavirus associated with human respiratory disease in China. Nature. 2020;579(7798):265–9.

7. Andersen KG, Rambaut A, Lipkin WI, Holmes EC, Garry RF. The proximal origin of SARS-CoV-2. Nature medicine. 2020;26(4):450–2.

8. Fehr AR, Perlman S. Coronaviruses: an overview of their replication and pathogenesis. Methods in molecular biology (Clifton, NJ). 2015;1282:1–23.

9. Wu A, Peng Y, Huang B, Ding X, Wang X, Niu P. Genome Composition and Divergence of the Novel Coronavirus (2019-nCoV) Originating in China. Cell host & microbe. 2020;27(3):325–8.

10. Huang Y, Yang C, Xu XF, Xu W, Liu SW. Structural and functional properties of SARS-CoV-2 spike protein: potential antivirus drug development for COVID-19. Acta pharmacologica Sinica. 2020;41(9):1141–9.

11. Bosch BJ, van der Zee R, de Haan CA, Rottier PJ. The coronavirus spike protein is a class I virus fusion protein: structural and functional characterization of the fusion core complex. Journal of virology. 2003;77(16):8801–11.

12. Xia S, Zhu Y, Liu M, Lan Q, Xu W, Wu Y. Fusion mechanism of 2019-nCoV and fusion inhibitors targeting HR1 domain in spike protein. Cellular & molecular immunology. 2020;17(7):765–7.

13. Lu J, du Plessis L, Liu Z, Hill V, Kang M, Lin H. Genomic Epidemiology of SARS-CoV-2 in Guangdong Province, China. Cell. 2020;181(5):997-1003.e9.

14. Seemann T, Lane CR, Sherry NL, Duchene S, Gonçalves da Silva A, Caly L. Tracking the COVID-19 pandemic in Australia using genomics. Nature communications. 2020;11(1):4376.

15. Deng X, Gu W, Federman S, du Plessis L, Pybus OG, Faria NR. Genomic surveillance reveals multiple introductions of SARS-CoV-2 into Northern California. Science. 2020;369(6503):582–7.

16. Condo J, Uwizihiwe JP, Nsanzimana S. Learn from Rwanda’s success in tackling COVID-19. Nature. 2020;581(7809):384.

17. Kumar S, Stecher G, Li M, Knyaz C, Tamura K. MEGA X: Molecular Evolutionary Genetics Analysis across Computing Platforms. Molecular biology and evolution. 2018;35(6):1547–9.

18. Nei M, Kumar S. Molecular evolution and phylogenetics: Oxford university press; 2000.

19. Cleemput S, Dumon W, Fonseca V, Abdool Karim W, Giovanetti M, Alcantara LC. Genome Detective Coronavirus Typing Tool for rapid identification and characterization of novel coronavirus genomes. Bioinformatics (Oxford, England). 2020;36(11):3552–5.

20. Mavian C, Marini S, Prosperi M, Salemi M. A Snapshot of SARS-CoV-2 Genome Availability up to April 2020 and its Implications: Data Analysis. JMIR public health and surveillance. 2020;6(2):e19170.

21. Chiara M, Horner D, Gissi C, Pesole G. Comparative genomics suggests limited variability and similar evolutionary patterns between major clades of SARS-CoV-2. bioRxiv; 2020.

22. Guan Q, Sadykov M, Mfarrej S, Hala S, Naeem R, Nugmanova R. A genetic barcode of SARS-CoV-2 for monitoring global distribution of different clades during the COVID-19 pandemic. International journal of infectious diseases : IJID : official publication of the International Society for Infectious Diseases. 2020;100:216–23.

23. Mercatelli D, Giorgi FM. Geographic and Genomic Distribution of SARS-CoV-2 Mutations. Frontiers in microbiology. 2020;11:1800.

24. Rambaut A, Holmes EC, O’Toole Á, Hill V, McCrone JT, Ruis C. A dynamic nomenclature proposal for SARS-CoV-2 lineages to assist genomic epidemiology. Nature microbiology. 2020;5(11):1403–7.

25. Farris JS. The retention index and the rescaled consistency index. Cladistics. 1989;5(4):417–9.

26. Phan T. Genetic diversity and evolution of SARS-CoV-2. Infection, genetics and evolution : journal of molecular epidemiology and evolutionary genetics in infectious diseases. 2020;81:104260.

27. Walls AC, Park YJ, Tortorici MA, Wall A, McGuire AT, Veesler D. Structure, Function, and Antigenicity of the SARS-CoV-2 Spike Glycoprotein. Cell. 2020;181(2):281-92.e6.

28. Koyama T, Platt D, Parida L. Variant analysis of SARS-CoV-2 genomes. Bulletin of the World Health Organization. 2020;98(7):495.

