## Supplementary figures and images for "Genomic diversity analysis of SARS-CoV-2 genomes in Rwanda"

### Supplementary figure S1

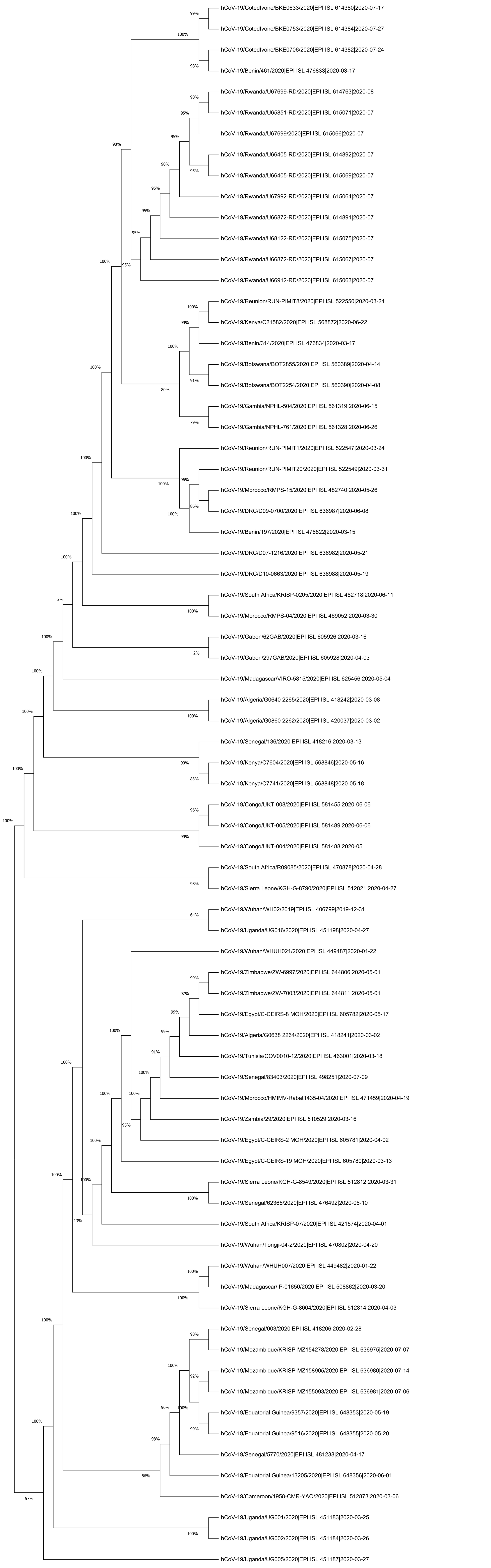
